# An organ-based multi-level model for glucose homeostasis: organ distributions, timing, and impact of blood flow

**DOI:** 10.1101/2020.10.21.344499

**Authors:** Tilda Herrgårdh, Hao Li, Elin Nyman, Gunnar Cedersund

**Affiliations:** Department of Biomedical Engineering, Linköping University, Linköping 58185, Sweden

**Keywords:** Glucose homeostasis, Glucose uptake, Insulin signaling, Mathematical modelling, Multi-level model

## Abstract

Glucose homeostasis is the tight control of glucose in the blood. This complex control is important and not yet sufficiently understood, due to its malfunction in serious diseases like diabetes. Due to the involvement of numerous organs and sub-systems, each with their own intra-cellular control, we have developed a multi-level mathematical model, for glucose homeostasis, which integrates a variety of data. Over the last 10 years, this model has been used to insert new insights from the intra-cellular level into the larger whole-body perspective. However, the original cell-organ-body translation has during these years never been updated, despite several critical shortcomings, which also have not been resolved by other modelling efforts. For this reason, we here present an updated multi-level model. This model provides a more accurate sub-division of how much glucose is being taken up by the different organs. Unlike the original model, we now also account for the different dynamics seen in the different organs. The new model also incorporates the central impact of blood flow on insulin-stimulated glucose uptake. Each new improvement is clear upon visual inspection, and they are also supported by statistical tests. The final multi-level model describes >300 data points in >40 time-series and dose-response curves, resulting from a large variety of perturbations, describing both intra-cellular processes, organ fluxes, and whole-body meal responses. We hope that this model will serve as an improved basis for future data integration, useful for research and drug developments within diabetes.

## Introduction

A dysfunctional glucose homeostasis is a hallmark for both type 1 and type 2 diabetes mellitus (T2D). In type 1 diabetes, the insulin-producing beta-cells are destroyed by the immune system. Since the other organs are unaffected, the treatment of T2D simply consists of insulin, taken via injections or insulin pumps. In T2D, the patient has both a reduced capacity to produce insulin and has developed a resistance to the hormone. This resistance appears in all of the three most metabolically active organs, which all respond to insulin: adipose tissue, muscle, and liver. Inside each of these organs, the response to insulin is governed by an interaction between intracellular signaling and metabolic networks. The resistance is spread between the organs, in ways which are not yet fully understood, but which involves numerous hormones, cytokines, and metabolites. To better understand this complex interaction, both in health and in disease, dynamic mathematical models are needed. Models for the top-level glucose homeostasis, involving a simple interaction between glucose and insulin, have been developed for decades ((Bergman et al. 1981). A first more advanced model (Dalla Man et al. 2007 was based on calculated flows of glucose and insulin between organs in response to a meal. A version of this model, trained on data from patients with type 1 diabetes, is approved by the Food and Drug Administration, FDA, for replacement of animal experiments in the approval of the algorithm inside new insulin pumps (Kovatchev et al. 2008. For more general applications, involving T2D, the intracellular insulin resistance must be combined with the whole-body interactions. Such models are called multi-level models.

There have been several efforts to create multi-level models of glucose homeostasis, reviewed in e.g. (Ajmera et al. 2013; Nyman et al. 2012; Nyman et al. 2016. One of the more comprehensive efforts is a series of nonlinear mixed effects models (Silber et al. 2007; Silber et al. 2010; Jauslin et al. 2007) developed to describe plasma levels of glucose and insulin after different interventions for single patients with T2D. Another effort has developed a glucose homeostasis model, based partly on (Dalla Man et al. 2007), to create a simulator to use in education and to simulate scenarios of disease (Maas et al. 2015). A third effort is the multi-level model of human glucose homeostasis we created 10 years ago (Nyman et al. 2011). This model contains the dynamic glucose-insulin interaction between organs in response to a meal, based on (Dalla Man et al. 2007). In this model, we sub-divided the original insulin-responding uptake in a muscle and a fat component, and linked the fat tissue glucose uptake to intracellular insulin signaling data, coming from our own studies. This link was possible since insulin-stimulated glucose uptake can be measured both in isolated adipocytes and in organs. The adipocyte uptake is measured *in vitro* together with insulin signaling data; the organ-level uptakes are measured using isotopic labelling and/or arteriovenous (AV) difference data, which measures the difference between arterial and venous blood. Since the uptake measurements from isolated adipocytes should correlate with the AV difference-based uptake-measurements for fat tissue, one can build a translation from *in vitro* to *in situ*, in humans. However, neither this model, nor any of the previously mentioned multi-level models, have subdivided the glucose uptake into the individual contributions of all of the main insulin-responding and glucose-utilizing organs: adipose tissue, muscle, and liver.

In this paper, we have updated the original multi-level connections in (Nyman et al. 2011), and resolved three critical questions or issues (Q1-Q3), regarding the role of each of the metabolically active organs in glucose uptake (Fig 1). More specifically, we have explicitly included the liver in the model as a glucose-utilizing organ, in contrast to the original models, which only considered it as a glucose producing organ (Q1). Secondly, we have included a timing difference between muscle and adipose tissue glucose uptake in the response to a meal (Q2). Thirdly, we have have updated the model to include the impact of blood flow on glucose uptake in adipose tissue (Q3). Finally, we merge these three improvements together with all of the other already published improvements described above, to an updated multi-level model (Q4). This model constitutes an updated view on the multi-level roles that each organ plays in glucose homeostasis, and allows for integration of future data for specific sub-systems into an integrated and more complete picture.

**Figure 1:**
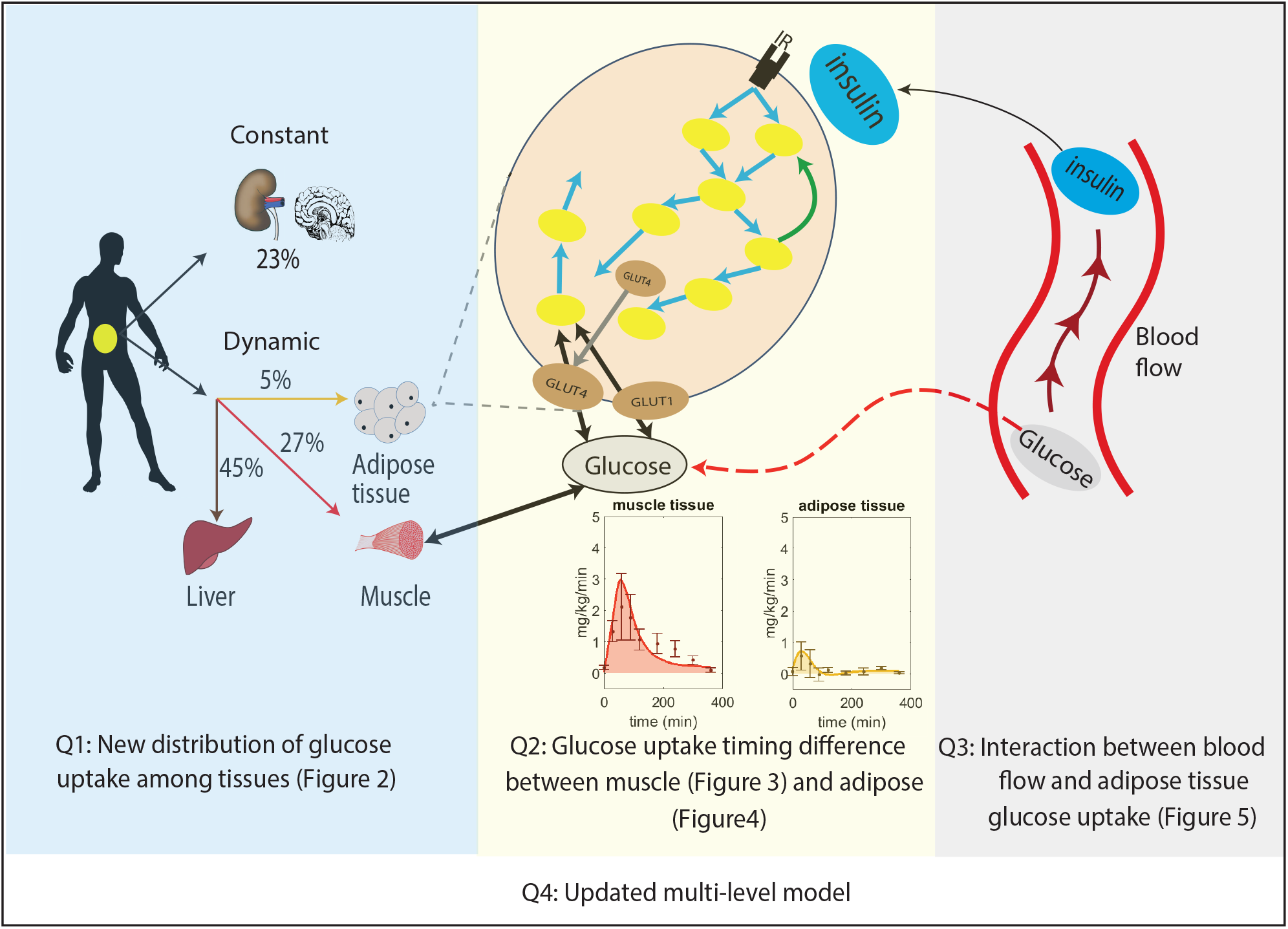
Overview of the improvements made to the original multi-level model. Q1: New distribution of postprandial glucose uptake among tissues in the whole-body level model. Q2: Implemented timing difference in glucose uptake between muscle and adipose tissue. Q3: Inclusion of the impact of blood flow on glucose uptake in adipose tissue. Q4: Merging Q1, Q2 and Q3 boxes together gives an updated multi-level metabolic model.

## Materials and methods

### Glucose dynamics in plasma and interstitial tissue

We have used ordinary differential equations (ODEs) in the standard form to build the models. All of the equations are given in the supplementary files, both as equations and as simulation files, and here we only describe the most central equations, relating to the changes done in this paper. The equations for the dynamics of the amount of glucose in interstitial tissue (*G*_*t*_) and plasma (*G*_*p*_) are given by

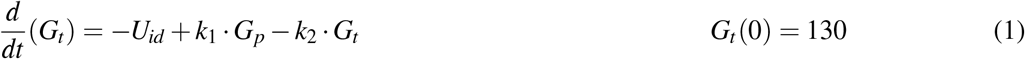

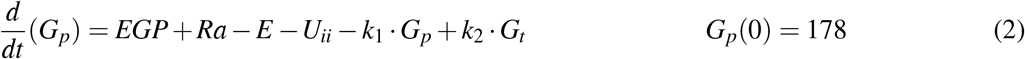

where *U*_*id*_ is insulin and glucose dependent glucose uptake, i.e. in fat, muscle, and liver; where *U*_*ii*_ is insulin independent and constant glucose utilization, i.e. glucose uptake by organs such as brain and kidneys; where *EGP* is endogenous glucose production from the liver; where *Ra* is glucose rate of appearance from the intestine; where *E* is glucose excretion through the kidneys; and where *k*_1_ · *G*_*p*_ and *k*_2_ · *G*_*t*_ denotes the flux from plasma to intestines and back, respectively. Note that *G*_*t*_ and *G*_*p*_ are states, while *U*_*id*_, *U*_*ii*_, *k*_1_ · *G*_*p*_, *k*_2_ · *G*_*t*_, *EGP*, *Ra*, and *E* are the reaction rates that describe flows of glucose. Similarly, *k*_1_ and *k*_2_ are parameters - rate constants - which are constant over time.

### Insulin-dependent and dynamic glucose uptake

The above equations are identical to those in the original Dalla Man model (Dalla Man et al. 2007), and the change that was implemented in (Nyman et al. 2011) was that *U*_*id*_ was sub-divided into a muscle and an adipose tissue part. We now sub-divide the insulin-dependent dynamic glucose uptake into three parts, i.e.

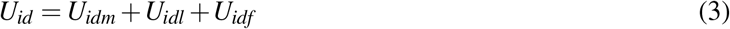

where *U*_*idm*_, *U*_*idl*_, and *U*_*idf*_ denotes the uptake rates into the muscle, liver, and fat, respectively. All of these uptake descriptions have changed to same extent, so let us now go through them one by one.

### Glucose uptake in muscle

Glucose uptake in muscle is given by

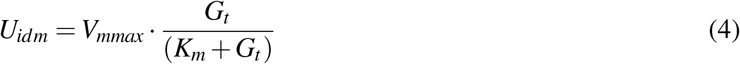

where *V*_*mmax*_ is the non-scaled maximal glucose uptake, and where *K*_*m*_ is the corresponding Michaelis-Menten parameter. The insulin-dependency of the glucose uptake is located in the expression for *V*_*mmax*_

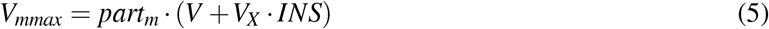

where *part*_*m*_ is a scaling parameter to balance the uptake of the muscle with the other organs, where *V* is the basal rate of glucose utilization, and *V*_*X*_ is the maximum rate of glucose entering the tissue (here muscle) from the surrounding tissue, and where *INS* denotes the interstitial insulin concentration. So far, these equations for the muscle uptake are the same as in (Nyman et al. 2011. In contrast, although *INS* is almost calculated in the same way as in (Nyman et al. 2011, the parameters describing the rate of entry and the rate of degradation are now allowed to be different, i.e.

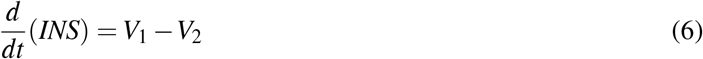

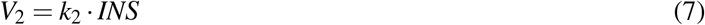

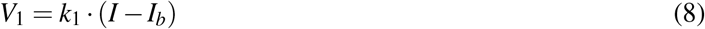

where *V*_1_ and *V*_2_ describe the rate of transport from the plasma and the rate of degradation, with corresponding rate constants *k*_1_ and *k*_2_, respectively; where *I*_*b*_ denotes the basal plasma insulin concentration; and where *I* denotes the insulin concentration in plasma.

### Glucose uptake in the liver

The liver was not included in the previous models, and thus its equations are new. They are similar to the equations for muscle, i.e.

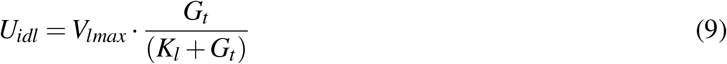

where *K*_*l*_ is a Michaelis-Menten constant, and and where *V*_*lmax*_ represents the maximum rate of glucose utilization in the liver. Just as for the equations for muscle, the insulin dependence is incorporated into the expression for *V*_*lmax*_, which is given by

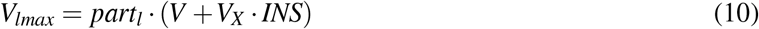

where *part*_*l*_ represent the relative glucose utilization of the liver in comparison with other tissues.

Note that the insulin-dependency of the liver glucose uptake is described as being direct, while in reality this dependency is indirect. Glucose uptake in the liver is done via the GLUT2 transporter, which is not regulated by insulin. In contrast, the glucose uptake in muscle and adipose tissue is done by the GLUT4 transporter, which is regulated by insulin. In the liver, insulin instead indirectly effects glucose uptake by up-regulating intracellular glucose phosphorylation and utilization. However, since the model is lacking intracellular reactions, this indirect effect present in the liver is approximated in the same way as the direct effect for the muscle. Note, finally, also that the EGP in the liver also is regulated by insulin, and that this is described as a separate process, in the same way as in (Nyman et al. 2011.

### Glucose uptake and metabolism in adipose tissue

Glucose uptake in the adipose tissue is the most advanced part of the model, since it is determined by intracellular processes, both regarding metabolism and regarding insulin signaling. The ultimate calculation of the uptake is given by the following expression

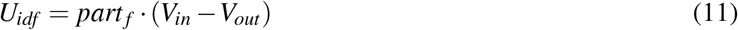

where *part*_*f*_ is a parameter, and where *V*_*in*_ and *V*_*out*_ describe the rate of glucose transport into, and out of, the cell, respectively. These two fluxes are given by

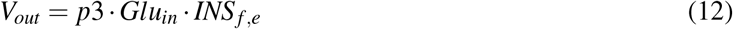

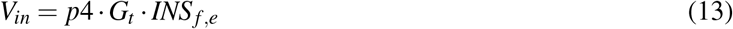

where *p*3 and *p*4 are transport parameters, where *Glu*_*in*_ is the amount of intracellular glucose, and where *INS*_*f,e*_ is the effect of insulin on these transport rates. These two equations show that the glucose uptake in the fat tissue depends on both intracellular metabolism, which alters the value of *Glu*_*in*_, and the intracellular signaling, which alters the value of *INS*_*f,e*_. The intracellular metabolism incorporates the first two steps of glycolysis, i.e. the steps involving intracellular glucose-6-phosphate (*G*_6*P*_). The equations are given by

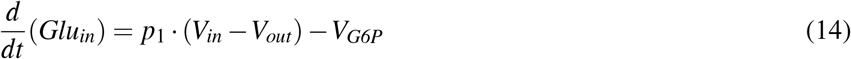

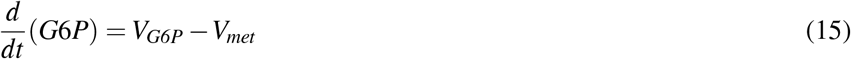

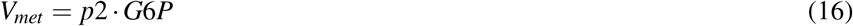

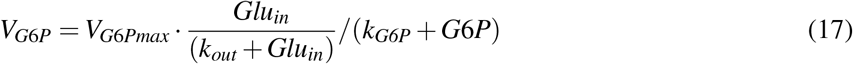

where *p*2, *p*1, *V*_*G6Pmax*_, and *k*_*out*_ are rate constants, and where *V*_*G6P*_ is the rate of phosphorylation of glucose. The intracellular insulin signaling is in itself the same as in (Brännmark et al. 2013, and it starts with insulin binding to the receptor (Supplementary eq 26), and ends with translocation of the GLUT4 transporter to the membrane (Supplementary eqs 46 and 45). What is new compared to (Nyman et al. 2011; Brännmark et al. 2013) instead concerns the usage of the GLUT4 transporter to calculate the resulting impact on glucose uptake, *INS*_*f,e*_. In our updated model, this insulin effect is given by

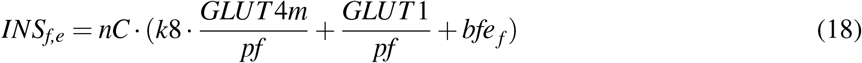

where *GLUT*4*m* is the amount of GLUT4 in the membrane, where *GLUT* 1 is the amount of GLUT1 in the membrane, and where *b f e*_*f*_ is the effect of blood flow; *nC*, *k*8, and *pf* are parameters. The GLUT4 and GLUT1 terms corresponds to the transport via the two glucose transporters, and *bfe*_*f*_ was introduced in (Nyman et al. 2011) as a scaling parameter between the data from the *in vitro* setting studying isolated adipocytes, and the *in situ* setting, where the adipose tissue is still located in the human body. In other words, the blood flow effect is not there when simulating *in vitro* experiments. In (Nyman et al. 2011), this difference in insulin effect was hypothesized to be dependent on blood flow, and in this paper, we show that such an impact on blood flow is indeed present. If one does not have data for the blood flow, the model will set *bfe*_*f*_ to a constant value, and if there is data for blood flow, we propose to use the new model described in the next section.

### Equations for the impact of blood flow on glucose uptake in adipose tissue

The impact of blood flow on glucose uptake is dependent on insulin. The same equations for adipose tissue is used for muscle glucose uptake (cf eqs (6)–(8)).

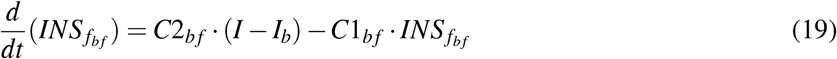

where *I* is insulin in plasma and *I*_*b*_ is the basal insulin level, and where *C*1_*bf*_ and *C*2_*bf*_ are rate constants.

Second, to calculate the impact of blood flow, we need to have an expression for how the blood flow is calculated. In this study, we only look at blood in controls, and in presence of Bradykinin, which increases the blood flow. This increase is also dependent on insulin. This control of blood flow, denoted *bf*_*f*_ is given by

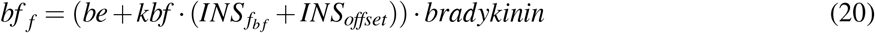

where *be* describes the direct effect of Bradykinin on blood flow; where *kbf* describes the combined effect of insulin and Bradykinin, and where *INS*_offset_ is a small offset introduced to make insulin concentrations positive (same as in (Nyman et al. 2011). The value of *bradykinin* is 1 in the absence of Bradykinin, and 30 in the presence of Bradykinin.

Finally, the blood flow and insulin are combined to impact the glucose uptake via the following expression for *bfe*_*f*_

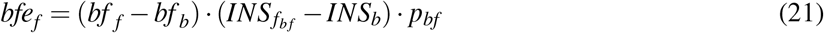

where *bf*_*b*_ is the basal blood flow, where *p*_*bf*_ is a paramete, and where *INS*_*b*_ is the basal insulin level in adipose tissue.

These are all the equations that have been changed in the current version of the model. The full set of ODEs from the final model, including the original simulation files, are found in the supplementary material. An interaction graph of the final model is given in Fig S1.

### Model simulation

Models were simulated in a modular fashion, by simulating part of the model with curves from other parts as input. Specifically, the new additions were simulated on their own together with equations for *G*_*p*_ and *G*_*t*_ with curves for *EGP*, *Ra*, *INS*, and *GLUT* 4*m*, simulated by the first model version presented herein (M1), as input.

### Parameter estimation

Parameter values for existing models are used from (Brännmark et al. 2013). The agreement between model simulations and experimental data is used to estimate values for new model parameters. This agreement is done by minimizing the distance between estimation data, denoted *y*, and corresponding simulated data for parameter *p*, which is denoted 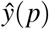. In our case, the estimation data consists of uptake rates of glucose into the adipose tissue and and muscle, which are denoted *U*_*idf*_ and *U*_*idm*_, respectively. The cost function used is the conventional weight least square, i.e.

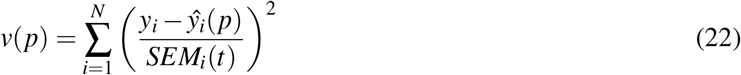

where the subscript *i* denotes the data point, where *N* denotes the number of data points, and where SEM denotes the standard error of the mean for the data uncertainty (Cedersund and Roll 2009).

We use a *χ*2-test to evaluate the agreement between model simulations and data. To be more specific, we use the inverse of the cumulative *χ*2 distribution function for setting a threshold, and then compare the cost function *v*(*p*) with a threshold.

In order to set that threshold, we need a significance level and the degrees of freedom. In this study, we use significance level 0.05, and the degrees of freedom used is specified for each analysis in the results.

Apart from the formal optimization described above, some additional ad hoc requirements were added to the parameter estimation. Specifically, to get a good estimate of the proportions of glucose taken up by the different tissues, a term adding a slightly increasing punishment for having a total uptake of glucose in liver higher than 50% or lower than 40% of total glucose uptake in all organs. The total glucose uptake of other organs except adipose tissue, muscle and liver (*U*_*ii*_) was punished in the same way for values higher than 28 % and lower than 18 % total glucose uptake of all organs.

The simple fitting to the impact of blood flow on glucose uptake was done by hand. A representative simulation was chosen for the comparison to the data uncertainties for total glucose uptake from Dalla Man (Dalla Man et al. 2007) (6).

### Uncertainty estimation

The uncertainty of the model simulations was estimated by, during the optimization process, saving all found parameters with an acceptable simulation according to section above.

### Code and data availability

We used MATLAB R2018b (MathWorks, Natick, MA) and the IQM toolbox (IntiQuan GmbH, Basel, Switzerland) for modeling. The experimental data as well as the complete code for data analysis and modeling are available at https://gitlab.liu.se/ISBgroup/projects/updated-multi-level.

### Experimental and clinical data

No new data were collected in this study. We therefore refer to the methods sections in the original articles (Frayn et al. 1993; Coppack et al. 1996; Gerich 2000; Moore et al. 2012; Iozzo et al. 2012; Brännmark et al. 2013) for the corresponding experimental methods.

## Results

### Distribution of postprandial glucose uptake between adipose, muscle, and liver (Q1)

The first improvement made to the original model (Nyman et al. 2011), referred to as M0, was to update the redistribution of the glucose uptake among the different tissues ((Fig 2A). The liver stands for almost half of the total postprandial glucose uptake (Fig 2B) (Gerich 2000), which was not explicitly accounted for in M0 (Fig 2A, dotted line). We therefore adopted the fluxes to fit to the data in 2B. More specifically, the liver was added as a glucose consuming organ, with a high net consumption compared to the other organs. In the updated model, referred to as M1 (Table S2), the liver is set to take up 45% of the total postprandial glucose uptake (Fig 2A-B), while adipose and muscle uptake were both reduced to 5% and 27% respectively. Furthermore, the glucose uptake by organs whose uptake is not affected by a meal (e.g. brain and kidneys) was increased to 23%. Note that in Fig 2A, this constant uptake is symbolized by the kidneys and the brain, because those are the most prominent glucose consumers (Gerich 2000), but that other tissues and organs can be seen as represented in this uptake as well.

**Figure 2:**
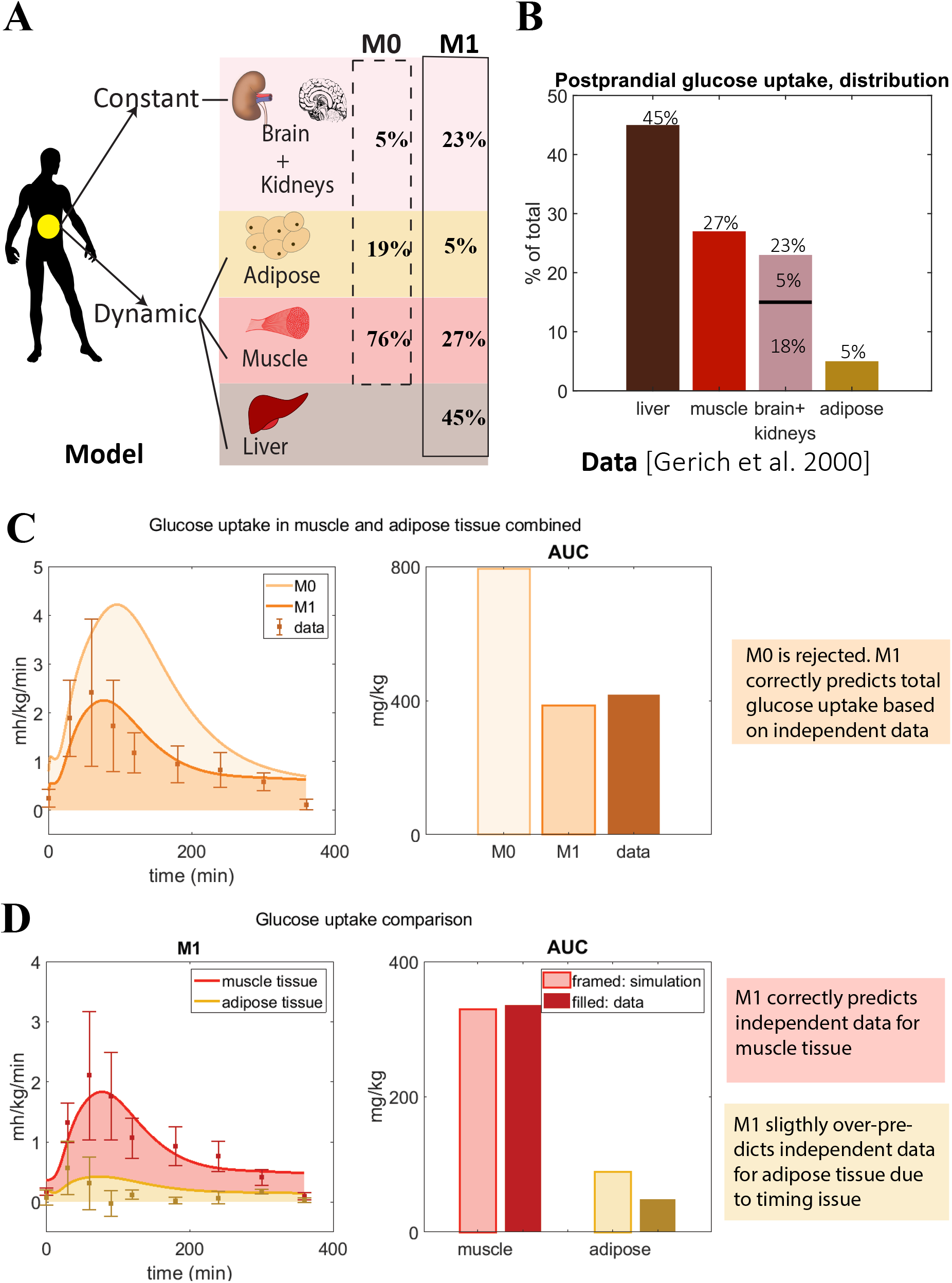
Updated distribution of glucose uptake among tissues. **(A)** In model M1, liver is added, the amount of glucose utilization in muscle and adipose is reduced, and the uptake that is constant during a meal of other tissues is increased compared to the original M0 model. **(B)** Glucose distribution among organs observed in data from (Gerich 2000). **(C)** Glucose uptake in muscle and adipose tissue combined for M1 and M0. The area under the curve for M0 is higher than seen in data from (Frayn et al. 1993; Coppack et al. 1996), and M0 is thus rejected. **(D)** Comparison between model M1’s predictions of adipose and muscle glucose uptake with new data not used for parameter estimation (Frayn et al. 1993; Coppack et al. 1996).

As a validation of these changes, we compared the resulting model simulations with data from other studies. More specifically, we compared the uptake of glucose in adipose and muscle tissue, as simulated by the two models M0 and M1, with data that measures the uptake in these two organs specifically. Such measurements are possible using e.g. AV difference data. In Fig 2C, the area under the curve (AUC) for M0 of adipose and muscle combined (dashed, light orange) is approximately 2 times bigger than the AUC of the data (solid, brown) in (Frayn et al. 1993; Coppack et al. 1996). This is clearly beyond the experimental uncertainty, and M0 is therefore rejected by a *χ*2 test 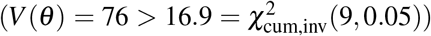. In contrast, M1 has approximately the same AUC as the data, and its simulations lies within the experimental uncertainty for most data points. Therefore, the time series is not rejected by the test based on these independent data 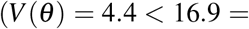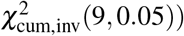. For these reasons, we reject M0, in favor of the new model M1.

A more detailed check of the quality of the updated model M1 is obtained by looking at the muscle and adipose tissue glucose uptake one by one (Fig 2D). For muscle (red), both the time-dynamics (left) and AUC (right) agrees between simulations (light red) and independent data (dark red). This visual observation is supported by a *χ*2 test 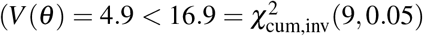. In contrast, the adipose tissue shows a reasonable agreement with data, but it is not quantitatively acceptable according to a *χ*2 test 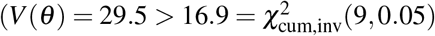. Looking closer at the time-series reveals that the value at the maximal uptake is fine, but that the problems lies in the fact that the dynamics of the uptake in muscle and adipose tissue are different, and that this is not captured in the model.

### Difference in time-resolved glucose uptake in adipose and muscle tissue (Q2)

Since the timing and agreement with dynamic glucose uptake in the muscle tissue is fine already in the model M1, this model was kept essentially intact. However, one minor modification that effects muscle uptake was introduced (Fig 3). In the previous model (M1), the rate constant of insulin transport into the interstitium (*V*_1_) is assumed to be the same (*k*_1_) as for the rate of the subsequent degradation of insulin (*V*_2_). Since there is no reason for these values to be the same, we updated the model to give these two reaction rates their own rate constants (*k*1, and *k*_2_, respectively). We refitted both parameters together with the other new parameters (introduced below) to the data, and the resulting model is referred to as M2a.

**Figure 3:**
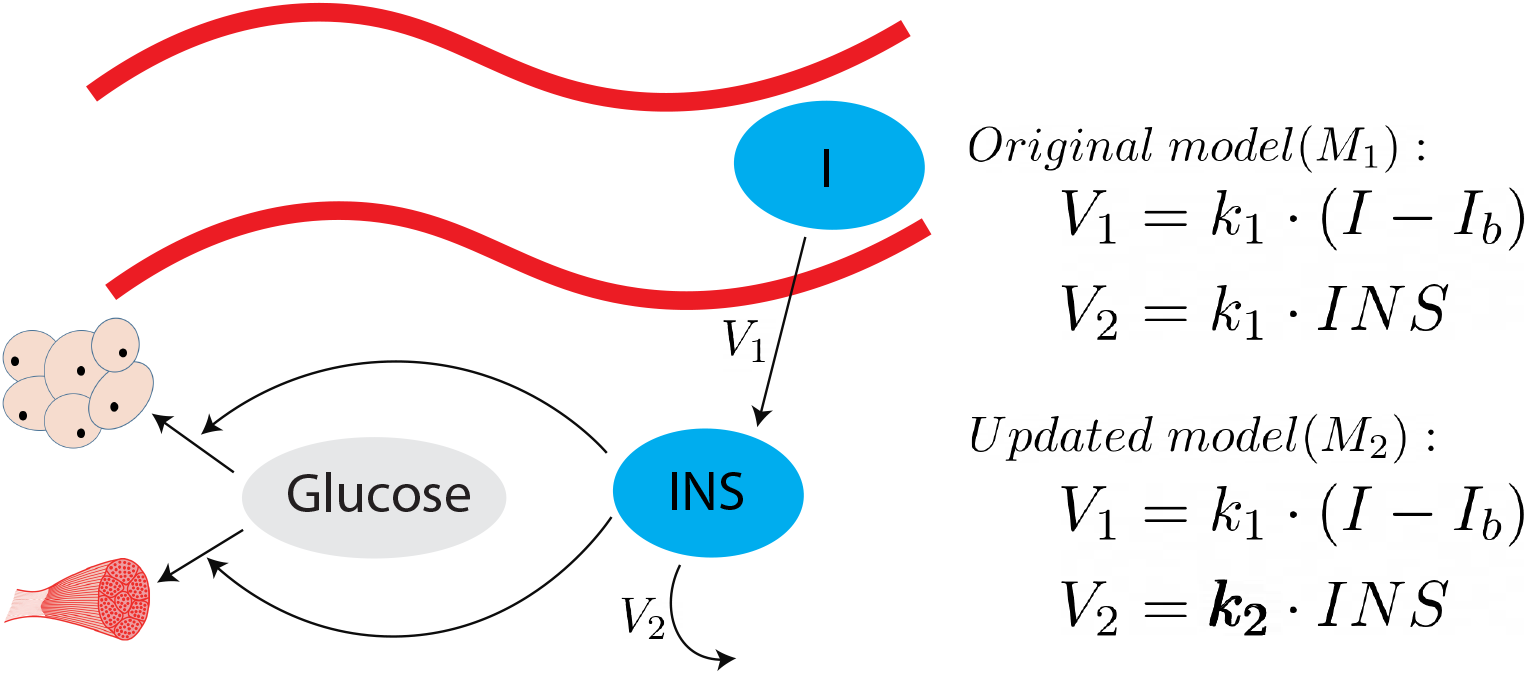
Illustration of insulin’s effect on glucose uptake, with the applied changes made in M2a. For M1, reaction V1 is dependent on parameter k1, while for M2, V2 is dependent on k2.

The developments for the adipose tissue glucose uptake needed to be more elaborate, and are available in Fig 4: the new model structure is depicted in Fig 4A and comparison with data is included in Fig 4B. As can be seen, the same difference as was introduced for muscle, M2a, yields a poor agreement with data for the adipose tissue, since the peak is too late. The main problem is that the glucose uptake in the adipose tissue has gone down to baseline levels already after around 100 min, when insulin levels still are high (Dalla Man et al. 2007) (7). Therefore, since the glucose uptake in the current model cannot go down before insulin goes down, an additional mechanism is needed. One such possible mechanism is the fact that the hexokinase reaction has a product inhibition (May and Mikulecky 1983). This leads to two new states in the next version of the model (M2b, 4A, red circle): intracellular glucose *GLU*_*in*_ and phosphorylated glucose, G6P. As seen, there is an inhibition from G6P to the rate of phosphorylation of *GLU*_*in*_.

**Figure 4:**
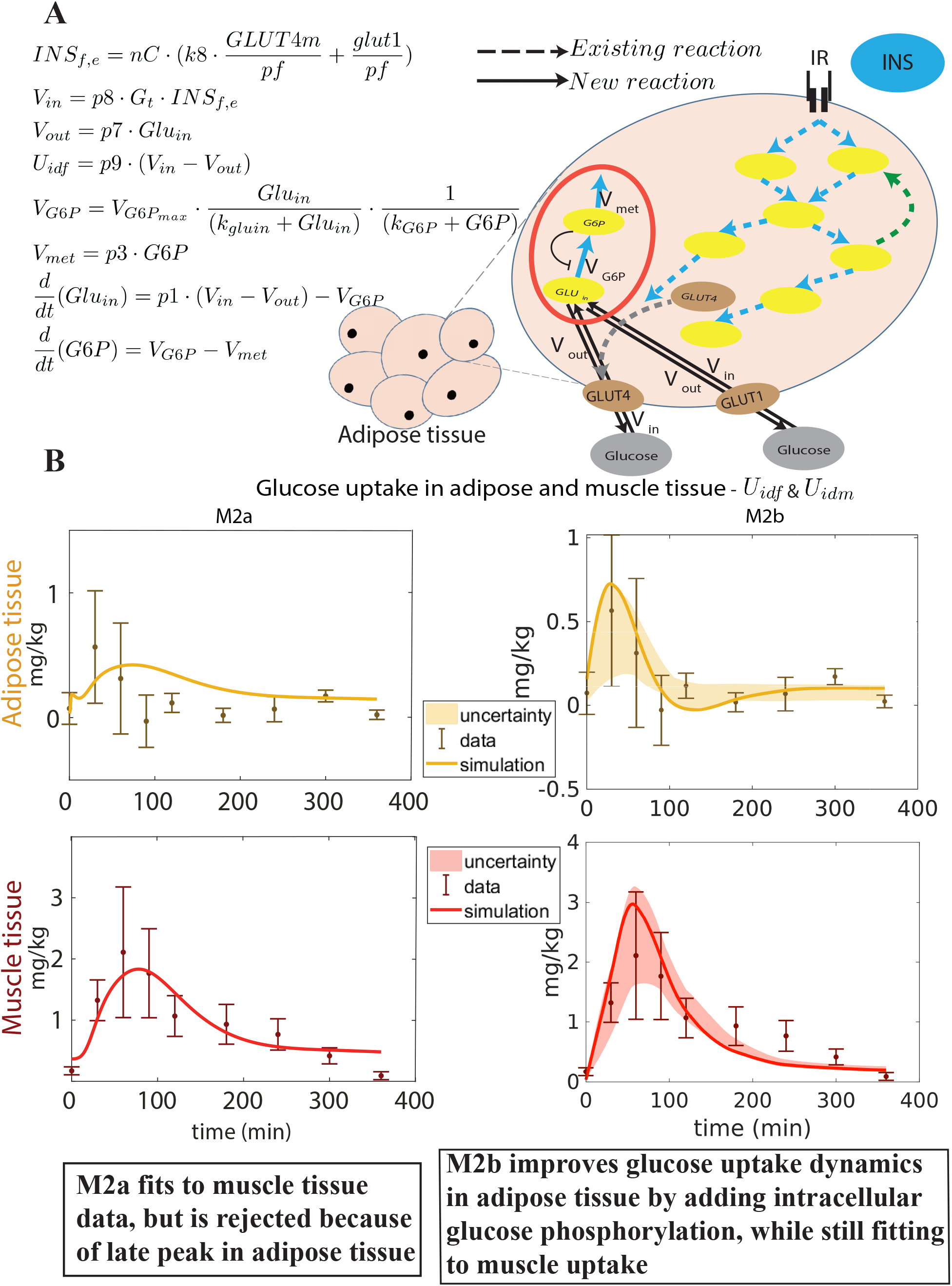
Improved dynamic behaviour of adipose tissue glucose uptake by improved intracellular module. **(A)** Illustration of the new intracellular adipose tissue module and ODE equations. The flow of glucose in to the cell, *V*_*in*_, is dependent on the amount of glucose in interstitium (*G*_*t*_) and inside the cell (*Glu*_*in*_), and the amount of *GLUT* 4*m* and *GLUT* 1 membrane glucose transporters through *INS*_*f,e*_. The out flow, *V*_*out*_, is only dependent on *Glu*_*in*_, which in turn depends on,*V*_*in*_, *V*_*out*_, and the phosphorylation of glucose into G6P (*V_G_*_6*P*_). The rate of phosphorylation to *G*6*P* is only dependent on *V*_*G6P*_ and the usage of *G*6*P* in metabolism (*V*_*met*_). **(B)** Timing comparison between uptake seen in data and the two models: M2a without phosphorylation, and M2b with glucose phosphorylation. In M2b, the peak comes earlier and the quantity of glucose taken up is closer to data than in M2a.

This modification allows for the following chain-of-events. When glucose uptake begins, the amount of intracellular glucose starts to build up, which is then phosphorylated into G6P. When the G6P reaches saturation levels, G6P inhibits the phosphorylation process from intracellular glucose, which leads to increasing intracellular glucose levels. Since the net glucose uptake is driven by the gradient across the cell membrane, this increase in intracellular glucose will decrease the glucose uptake, even though insulin levels still might be high. The resulting simulations of glucose uptake in muscle and adipose tissue (Fig 4B, right), agrees with the data both according to a visual check, and according to a *χ*2 test 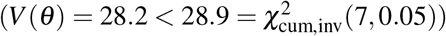.

### Improvements in the intracellular adipose tissue model: glucose metabolism and blood flow effects (Q3)

The final improvement made was the addition of the impact of blood flow on insulin-stimulated glucose uptake in the adipose tissue. This interaction was hypothesised in Iozzo *et al*, where they looked at the effect of blood flow and insulin, separately and combined, on glucose uptake in adipose tissue (Iozzo et al. 2012) (Fig 5A-B). Increased blood flow was achieved with the drug Bradykinin. In these experiments, Iozzo *et al* observed that glucose clearance was not significantly changed when only adding Bradykinin (Fig 5B, left). In contrast, when combining both Bradykinin and insulin, the glucose uptake is increased compared with only adding insulin (Fig. 5B, right). The same behaviour is produced by the model in Fig 5C, where the glucose uptake only increases when both Bradykinin and insulin is present. The parameter *bradykinin* was changed from 1 to 30 to represent the addition of Bradykinin, and the parameter *INS*_offset_ is changed from 0 to 7 to represent insulin infusion (Fig 5A). This behaviour also agrees with data according to according to a *χ*2 test 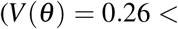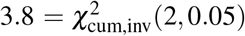, where the degrees of freedom have been compensated for with the number of new parameters, 4-2=2). The updated model is referred to as M3, and as for the other model additions, the new equations are shortly depicted in the figure (here Fig 5A), and described in detail in Materials and Methods and Supplementary files.

**Figure 5:**
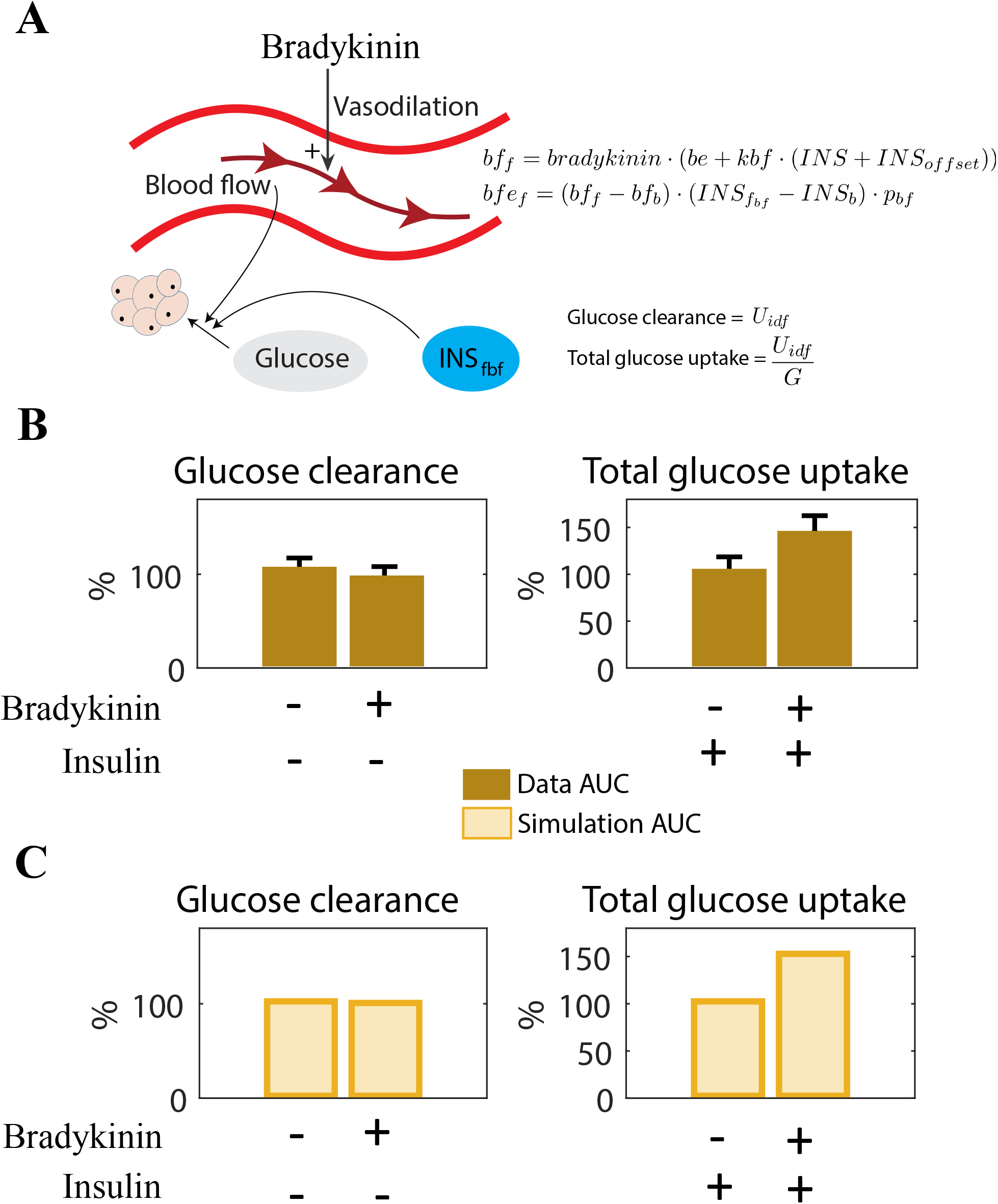
Interaction between blood flow and insulin on adipose glucose uptake. **(A)** Illustration of blood flow and insulins effect on adipose tissue glucose uptake. The drug Bradykinins increases the blood flow. The new equations for blood flow (*bf*), dependent on bradykinin (*BRF*), and blood flow effect (*b f e*_*f*_), dependent on blood flow dependent insulin in fat tissue (*INS*_*fbf*_) **(B)** Behaviour seen in data as response to insulin and Bradykinin. Insulin alone has a relatively small effect on glucose clearance, but increases glucose uptake significantly when combined with Bradykinin (Iozzo et al. 2012). **(C)** The same behaviour as in **(B)** (Iozzo et al. 2012) can be simulated with the model. Adding Bradykinin is simulated by increasing the value of *bradykinin*, and adding insulin infusion is simulated by increasing the value of *INS*_offset_ from 0.

### The final model (Q4)

Finally, we consider the performance of the resulting final multi-level model, in relation to all of the data that has been generated over the years. The final model can fit to dynamic data of postprandial glucose uptake in both adipose and muscle tissue (Fig 6A, same data as in Fig 2D, from (Coppack et al. 1996). The same figure displays predictions of dynamic uptake in the liver (for which the same type of AV difference data is non-existent), and for the tissues with a constant demand of glucose (such as the brain). Finally, the right-most sub-figure in Fig 6A shows that the model agrees well with the total dynamic glucose uptake from (Dalla Man et al. 2007).

**Figure 6:**
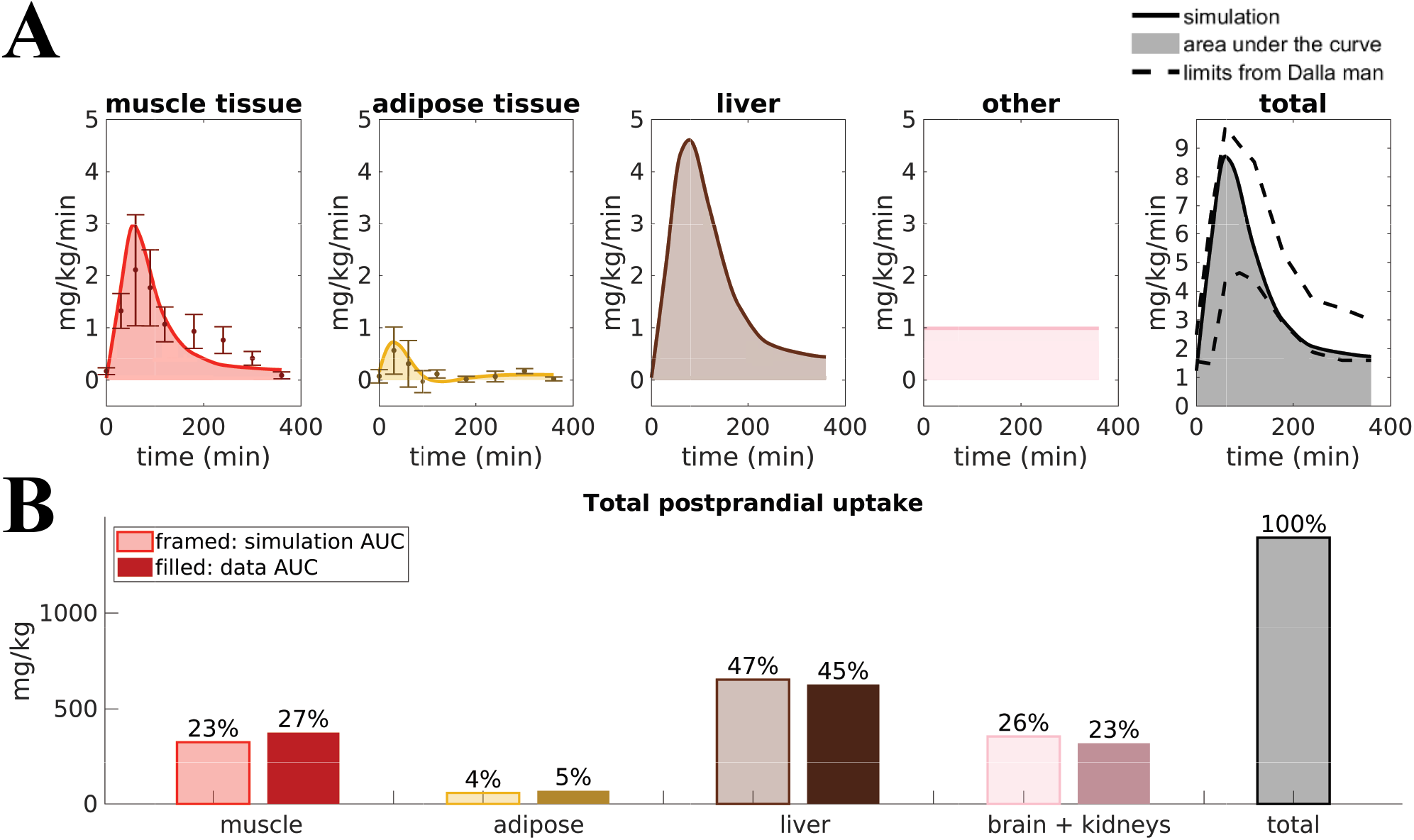
The behaviour of the final updated multi-level model. **(A)** Simulations of glucose uptake in all organs and tissues and time-series for the data used to fit the model(Coppack et al. 1996). The total glucose uptake is within the bounds presented in (Dalla Man et al. 2007). **(B)** Total glucose uptake for all organs, simulated by the final model and from the data used to fit the model (Coppack et al. 1996).

Furthermore, the AUCs for the different tissues in the final combined model is in line with the corresponding AUC data (Fig 6B), just as they were in step Q1 (Fig 2B). The two left-most bars, for muscle and adipose tissue, are given by the AUC of the corresponding time-series in Fig 6A (cf Fig 2D), and the liver and brain/kidney uptake are the same as in Fig 2B.

The final model is also in agreement with data previously used in the model development. The agreement with the most important such data sets are re-plotted in Fig 7 (Dalla Man et al. 2007), which describes meal responses for the following variables: Plasma Glucose, Plasma insulin, Endogenous Glucose Production, Glucose Rate of Appearance from the intestines, Glucose uptake or utilization, and insulin secretion. As can be seen, the model simulations (lines) are within the experimental uncertainty (grey area) for all these time curves (agreements between simulation and data are similar as in (Dalla Man et al. 2007).

**Figure 7:**
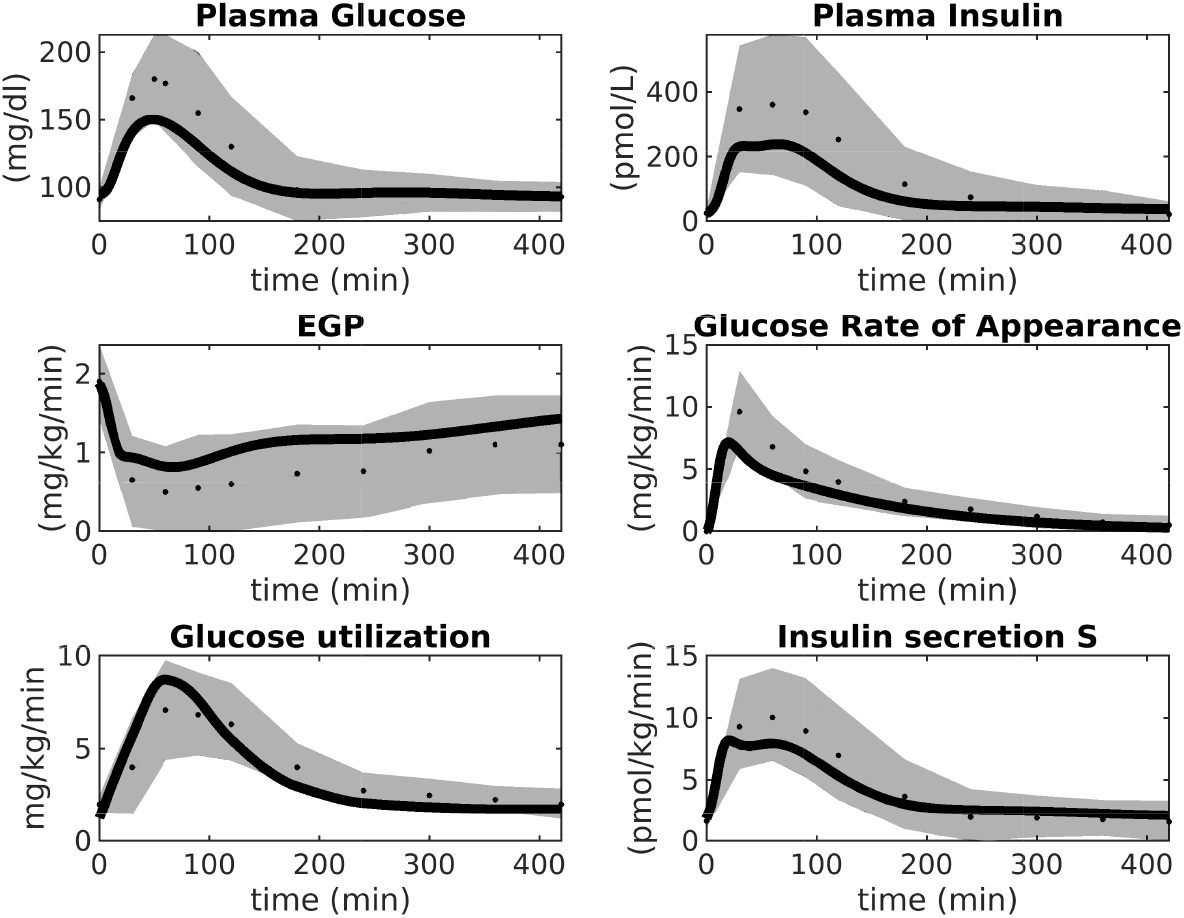
Simulations of M4 (lines) in comparison with data (dots) and uncertainties (grey areas) from (Dalla Man et al. 2007). M4 simulations are within the data uncertainties established in(Dalla Man et al. 2007)

Similarly, because of the hierarchical way that the multi-level model is constructed, it also still agrees with all of the intracellular signalling data, which we have collected over the years (Brännmark et al. 2013). The most important such data is depicted in Fig 8. These data (error bars) describe time-series and dose-response curves in response to insulin for a number of intracellular proteins: the insulin receptor (IR), the insulin receptor substrate-1 (IRS1), protein kinase-B (PKB), Akt-substrate 160 (AS160), Ribosomal protein S6 kinase beta-1 (S6K1), Ribosomal protein S6 (S6), as well as cellular glucose uptake. The model simulations (lines) are in agreement with both data from non-diabetic and lean controls (blue), and from obese people with type 2 diabetes (red), with changes only in a few key parameters (for more details, see (Brännmark et al. 2013). Similar agreements for additional proteins - such as extracellular signal-regulated kinases (Erk1), ETS Like-1 protein Elk-1 (Elk1), Forkhead box protein O1 (FOXO1), etc - is equally possible to obtain by replacing the intracellular part of the model with those in (Nyman et al. 2014; Rajan et al. 2016).

**Figure 8:**
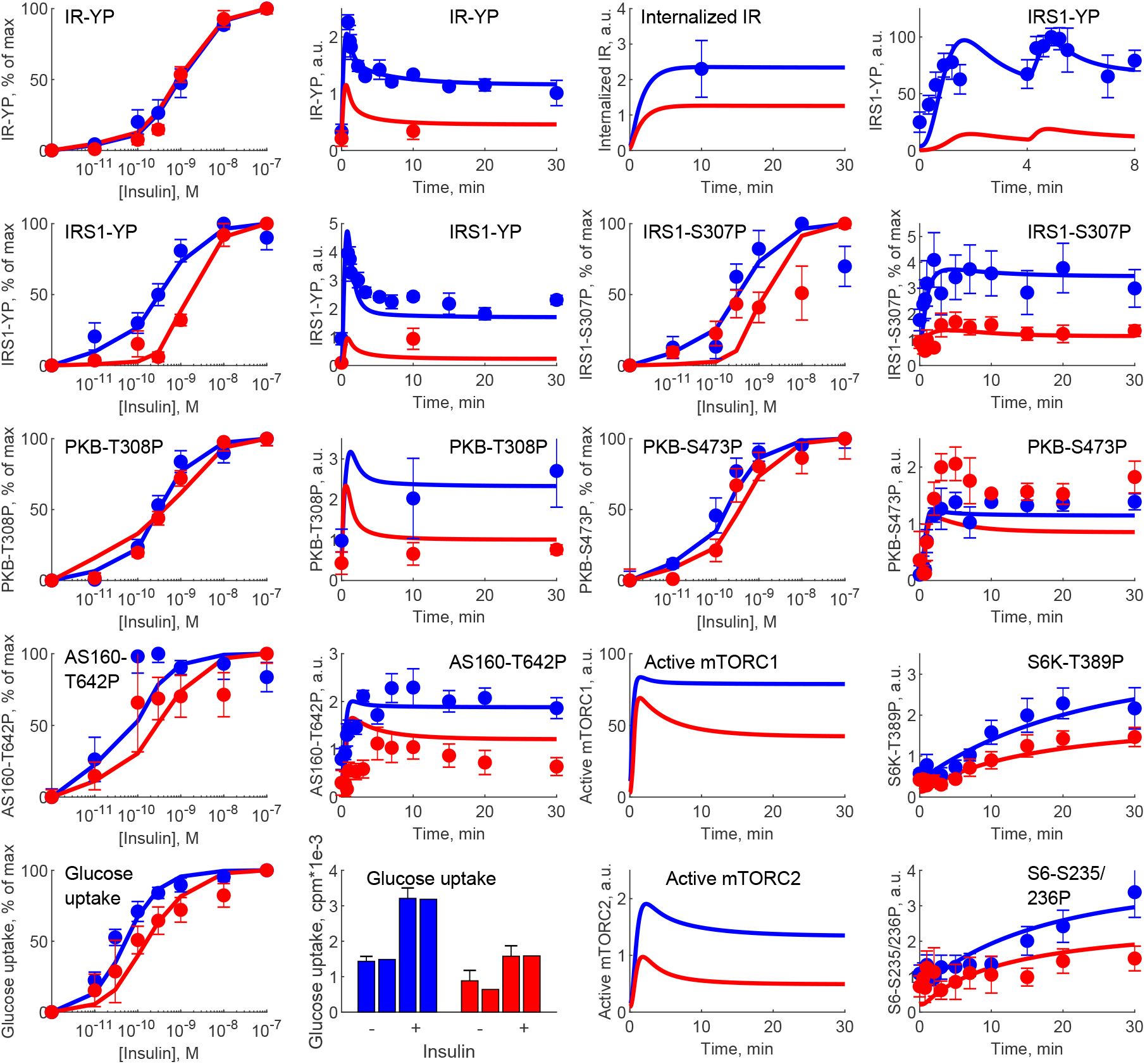
Simulations of M4 (lines) in comparison with experimental data (error bars) from (Brännmark et al. 2013). M4 can describe data for intracellular insulin signaling in adipocytes, both normally (blue) and in T2D (red). IR, insulin receptor; IRS1, insulin receptor substrate-1; PKB, protein kinase-B; AS160, Akt-substrate 160; S6K1, Ribosomal protein S6 kinase beta-1; S6, Ribosomal protein S6; YP, tyrosine phosphorylation; SP,serine phosphorylations (on sites 235/236, 307, 473); TP, threonine phosphorylation (on site 308).

## Discussion

Glucose homeostasis is a complex multi-organ and multi-level system, which requires multi-level mathematical modelling for a full understanding. We have herein improved an existing such model (Nyman et al. 2011) for glucose fluxes in the circulation, linked to intracellular pathways in adipocytes, in response to a meal. Specifically, we have (Q1) made a new subdivision of glucose uptake between all relevant organs, to provide more reliable proportions and to include uptake in the liver (Fig 2); (Q2) improved the elimination of interstitial insulin to be tissue-specific (Fig 3), and included intracellular metabolism of glucose inside adipocytes, to capture an earlier peak in the glucose uptake in adipocytes compared to the corresponding peak in plasma insulin (Fig 4); and (Q3) accounted for the impact of blood flow on glucose uptake (Fig 5). The final combined model (Q4) can fit to all of the new data for glucose uptake in all organs (Fig 6), as well as to all previous data, such as the postprandial glucose and insulin fluxes and concentrations in (Dalla Man et al. 2007) (Fig 7), and the intracellular data in (Brännmark et al. 2013) (Fig 8). To the best of our knowledge, this is the most comprehensive description of such a wide variety of data for glucose homeostasis in humans, and we hope that it will become a useful resource also for integration of future data.

One of the main contributions in this work is the addition of glucose uptake in the liver (Q1). This addition is important because the liver is the the organ that takes up the most glucose: approximately 45% (Fig 2). Apart from this, the liver has a unique function in glucose homeostasis, since it is the only organ that can produce glucose from other metabolites. These two functions, glucose uptake and endogenous glucose production (EGP), are now modeled as separate processes. In other words, the liver can both produce and take up glucose at the same time. While there may be situations when only the net uptake/release is important, there are also situations when one can experimentally resolve the two fluxes. For instance, when labeled metabolites have been ingested, one can see the rate by which these are converted to glucose and secreted, even in postprandial conditions, when the net effect of glucose transport is into the cell. Such data have previously been used to train the EGP fluxes (Fig 7) (Dalla Man et al. 2007), and we have now added corresponding data for glucose uptake (Fig 6B). Note that this model is only fitted to the data in Fig 2B, and that the agreements seen in Fig 2C-D serve as a simple validation of this part of the model. With this said, it should be emphasized that both the muscle and the new liver module are highly simplified. Only the muscle and adipose modules have been tested with respect to dynamic uptake data, and only the adipose module with an intracellular signaling part, based on detailed intracellular data, resolving the complicated intracellular metabolic fluxes. These limitations are present primarily because such data are rare or non-existent.

At the heart of resolving both Q1 and Q2 lies measurements of glucose fluxes, which have been measured in a variety of ways. The glucose fluxes from (Dalla Man et al. 2007) was based on a triple tracer protocol, which allows for the simultaneous calculation of plasma glucose, EGP, glucose rate off appearance, and glucose utilization (Fig 7). These data are based on advanced calculations, which in turn are based on various assumptions and mathematical models developed within the field of tracer based measurements (Wolfe et al. 2005). These particular assumptions are not necessary in the organ specific glucose utilization curves, available e.g. for muscle and adipose tissue (Fig 2C and D). These data are based on an AV difference-based protocol, which samples in both an artery and veins that have past through either muscle or adipose tissue, and by looking at the difference between the ongoing and the outgoing blood (Coppack et al. 1996). This is a more direct way of measuring how each organs contributes to the glucose disappearance from the blood. Nevertheless, also AV-difference data does not measure glucose uptake in the primary cells, myocytes and adipocytes, respectively. This means that the quick decline in glucose uptake in adipose tissue (Fig 4B) could in fact be the result of a quick equilibrium between interstitial and capillary glucose concentration. One could possibly develop an alternative model based on that equilibration-based assumption, to explain the quick decline of the glucose uptake in the adipose tissue, either as a replacement or as a complement to the herein implemented mechanism based on product inhibition (Fig 4A). Finally, the fact that the model is based on three different types of measurements of glucose uptake (cellular *in vitro*, tracer-based, and AV-difference based), and can describe all of these types of data simultaneously, is a reason why a relatively simple validation, such as that in Fig 2C-D, still is of value.

The final question addressed herein (Q3) concerns the impact of blood flow on glucose uptake, which is highly simplified because the real relationship is a bidirectional one. The data in Fig 5B shows that glucose uptake is increased by increased blood flow, at least in cases when insulin is present. This relationship is captured in the final model. However, that model can only describe situations where the blood flow is altered in a way that is not connected to the metabolic response, such as when adding Bradykinin (Fig 5B). In other words, the model cannot describe meal-induced blood flow changes and its associated impact on glucose uptake. The development of a model for blood-flow regulation during e.g. meal-responses is an important task for future modelling works. Another weakness regarding the blood flow part of the model concerns the lack of validation. The model is only fitted to the data in Fig 6B. In the analysis, we compensate for that by reducing the degrees of freedom from the number of data points (4) to the number of data points minus the number of parameters (4-2=2). However, one could argue that the two baseline bars should not be counted since they are normalized to be 100%. In such an interpretation, the degrees of freedom are 0, a chi2 test can not be done, and the only possible assessment of the quality of the model is a visual comparison of the differences between Fig 5B and C. For all these reasons, the blood flow part of the model is to be considered as a first step in the development of a model for the blood flow and its function in glucose homeostasis.

It is important to compare the model presented herein to other similar models in the literature. In the introduction, we mentioned the now classical nonlinear mixed effects models describing plasma levels of glucose and insulin (Silber et al. 2007; Silber et al. 2010; Jauslin et al. 2007). These models have since these early publications been used to scale data between pre-clinical data from animals to clinical human data for glucose and insulin concentrations (Alskär et al. 2017), and to describe cross-talk with more long-term processes, such as disease development in mice (Choy et al. 2016) and dynamics of HbA1c (Kjellsson et al. 2013; Møller et al. 2013). Glucose homeostasis-centered models, focusing on the glucose-insulin interplay, lie at the heart of mathematical models developed for type 1 diabetes, e.g. to aid insulin-pumps, and to develop a so-called artifical pancreas(Huang et al. 2012; Fabris and Kovatchev 2020). Another application of glucose homeostasis models exist for meal response T2D simulator model, developed for pedagogical and motivational purposes (Maas et al. 2015). None of these models have subdivided glucose uptake in the different organs, or included intracellular responses, in multi-level and multi-organ models. There exists one model that does this, developed by Uluseker *et al* (Uluseker et al. 2018). This multi-level model is based on a version of the Dalla Man model (Dalla Man et al. 2007) connected with our intracellular adipocyte model (Brännmark et al. 2013), while also including hormonal effects on glucose intake/appetite (leptin, ghrelin) and insulin levels (incretin). However, this model does not compare their whole-body simulations with any data, and does not include the liver as a glucose consuming organ.

The model presented in this work only included intake of glucose, and thus discarded the effects of proteins and fat on the meal response, something that other models do take into account, to some extent. Sips *et al* developed a model that integrates fatty acids with glucose metabolism (Sips et al. 2015), but this model needs a triglyceride curve as input, and lacks protein metabolism. Nevertheless, the Sips model is another expansion of the Dalla Man model (Dalla Man et al. 2007) and can thus be merged with the developments herein. Two models that include protein and fat intake from a meal are the ones developed by Hall *et al* and Sarkar *et al* (Hall et al. 2011; Sarkar et al. 2018). These models are however developed for long term simulations (over several years), and can thus not simulate a meal response. In similarity to the model presented here, the Sarkar model include liver, muscle, and adipose tissue as glucose consuming organs, but in contrast also adds the pancreas as a glucose consuming organ. Furthermore, the Sarkar model disregards the organs taking up a constant amount of glucose (brain and kidneys). In any case, the Sarkar model only describes data for long-term dynamics, and does not describe meal-respones. Another longitudinal model describing glucose dynamics on both short and long-term time-scale is the one developed by Ha *et al*. (Ha and Sherman 2019). This model is, in contrast to the other two longitudinal models mentioned above, multi-scale in that it can look at both changes over years, including the progression towards diabetes in a semi-mechanistic fashion, as well as meal response dynamics happening in the scale of hours and minutes. This model does, however, not include the distribution of glucose among different organs.

There are also some multi-level and multi-scale models for other systems that should be mentioned. One such model is the one developed by Barbiero *et al* (Barbiero and Lió 2020). This model combines whole-body dynamics with the function of organs and individual cells, and is able to simulate dynamics in seconds up to several days. The model was used to simulate the cardiovascular and inflammatory effect of both T2D diabetes and COVID-19, using personalized parameters. However, this model has an important short-coming: its simulations are not compared with any data. There also exists interconnected models for e.g. heart function, describing the function of cardiac cells up to the integrated behavior of the intact heart (Smith et al. 2009).

In summary, there does not exist any other multi-level model describing glucose meal response, that also separates between the different organs’ glucose uptake. In this work, we present such a model, that, due to its modular approach can be easily expanded in different directions. This expansion-possibility is due to both the modular structure, and to the fact that each module can be treated as a separate modelling problem. In other words, as long as the model for each module agrees with the input-output profiles of insulin and glucose, the new model can replace the old model, with little alterations on whole-body dynamics. In the earlier developed model (Nyman et al. 2011), we took this modularity one step further, by replacing the simpler 5-state insulin receptor module with a much more detailed 37-state module for the receptor dynamics, including the possibility for a receptor to bind up to three insulin molecules (Kiselyov et al. 2009). This demonstrates the usefulness of developing a model in modules, so that the right level of details can be included depending on the data/questions you want to analyze.

Since the original publication of our first multi-level model (Nyman et al. 2011), we have built further on this model in several directions, and all of these developments can be re-used also in our new model. We have e.g. expanded the intracellular part to explain a more and more comprehensive picture of the alterations in intracellular signaling that occur in T2D. This has been done by taking adipose tissue biopsies from both healthy and T2D individuals, and characterise their respective insulin signalling. In (Brännmark et al. 2013), we presented a first model of how the insulin resistance occurs, and in subsequent works, we have added additional proteins, such as FOXO1 transcription factor (Rajan et al. 2016), insulin control of MAPKs ERK1/2 (Nyman et al. 2014). Because of the modular way that our multi-level model is structured, one can replace the herein used intracellular model with any of these other alternatives. The same expansions can be done also for other organs. We therefore hope that this multi-level model in the future can serve as a hub for connecting data and models together into a useful systems-level understanding.

## List of abbreviations

T2D: Type 2 diabetes
AUC: Area under the curve
EGP: Endogenous glucose production
AV: Arteriovenous
ODEs: Ordinary differential equations

## Acknowledgements

We thank the Swedish research council (2018-05418, 2018-03319, 2019-03767), CENIIT (15.09, 20.08), the Heart and Lung Foundation, the Swedish foundation for strategic research (ITM17-0245), SciLifeLab/KAW national COVID-19 research program project grant (2020.0182), H2020 (PRECISE4Q, 777107), The Swedish Fund for Research without Animal Experiments, and ELLIIT. This manuscript has been submitted to bioaRxiv.

## Supplemental Material

All the ODEs for the final model M4:

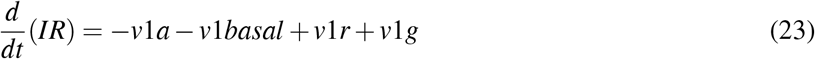

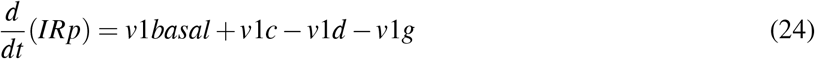

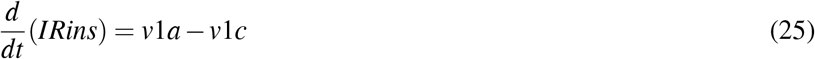

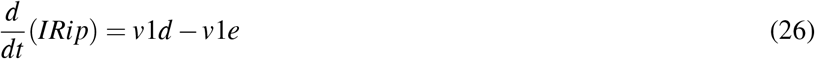

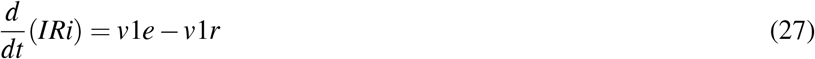

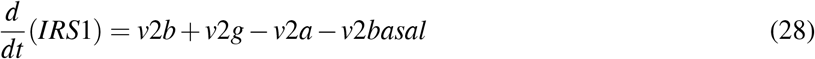

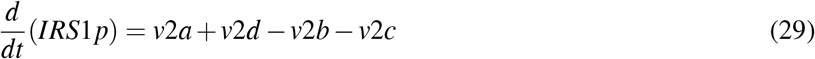

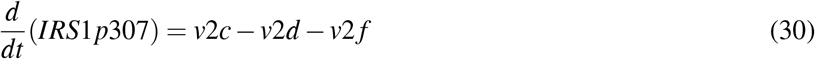

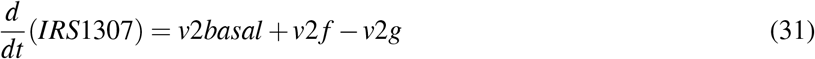

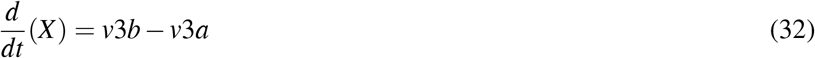

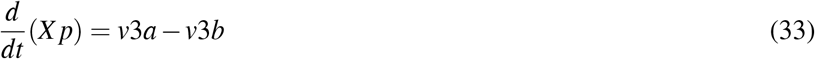

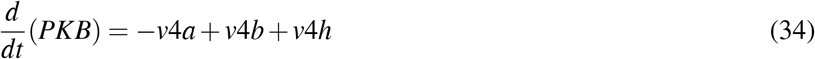

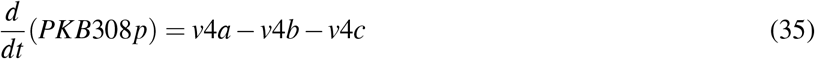

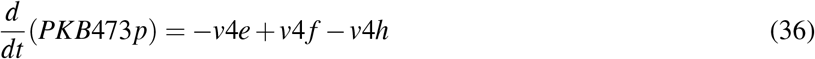

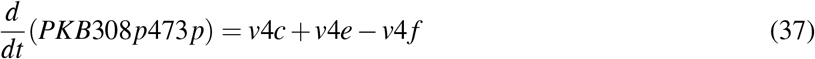

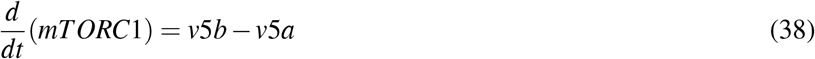

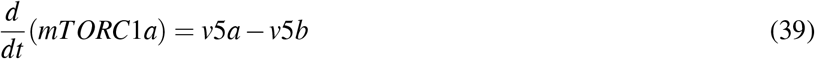

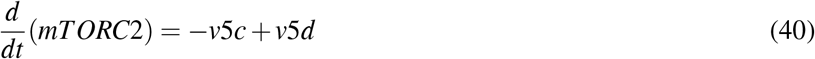

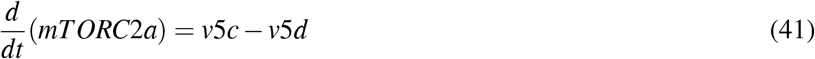

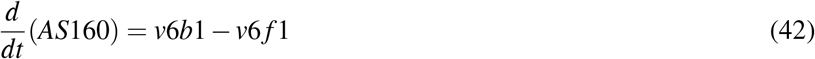

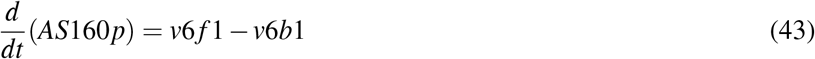

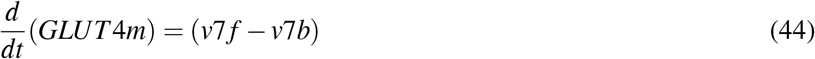

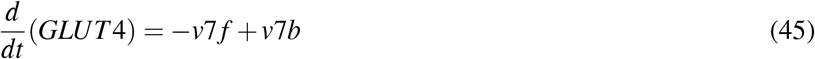

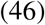

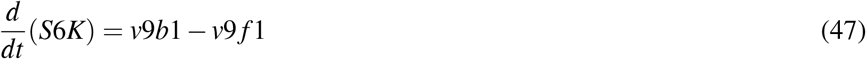

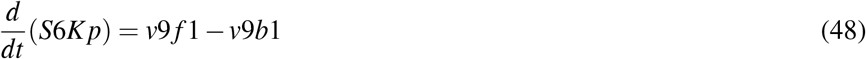

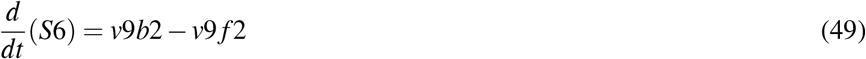

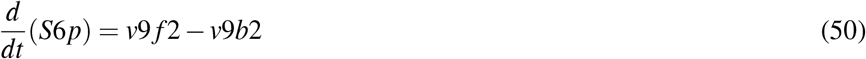

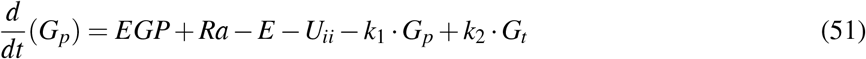

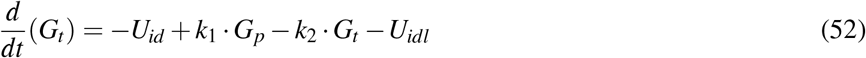

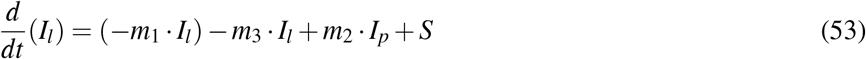

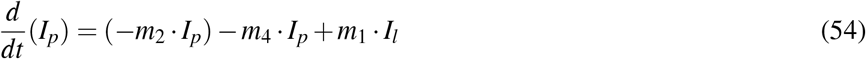

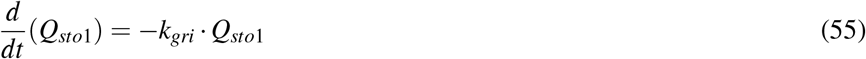

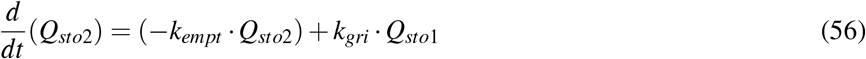

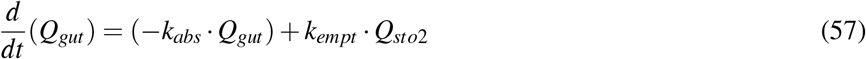

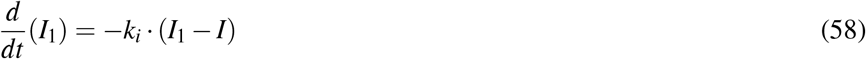

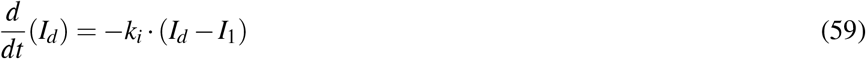

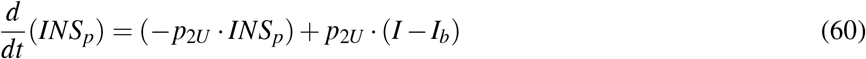

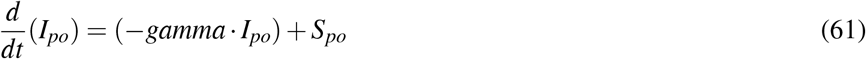

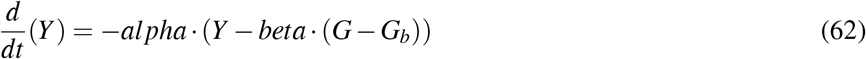

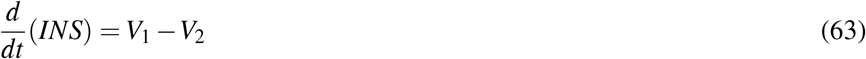

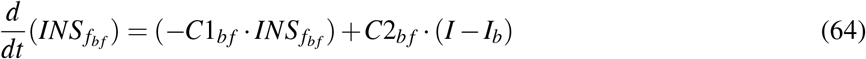

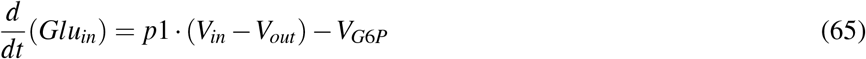

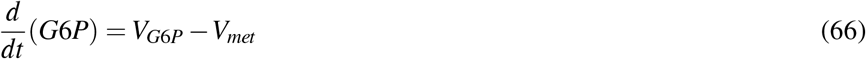

All variables of final model:

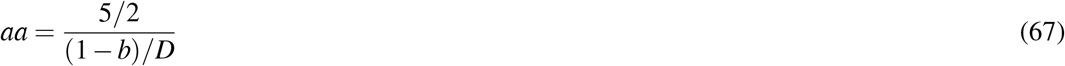

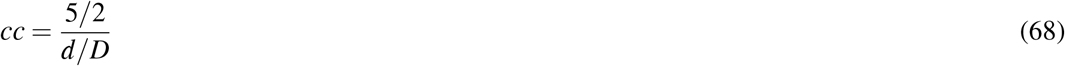

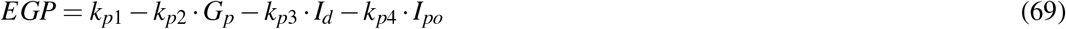

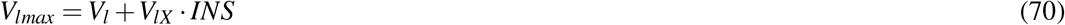

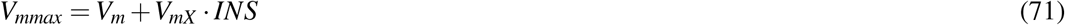

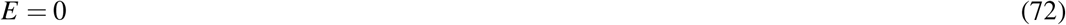

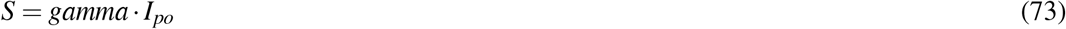

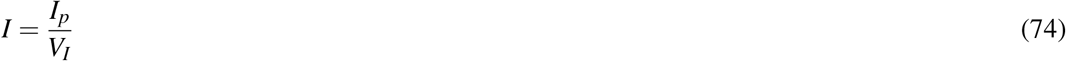

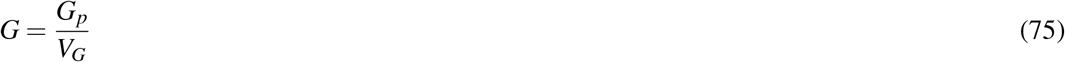

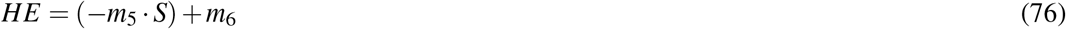

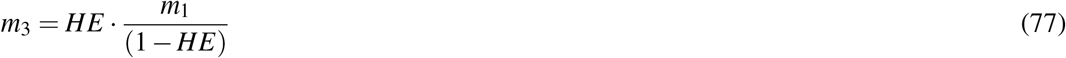

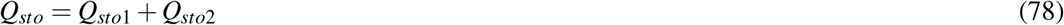

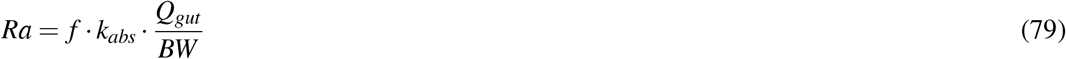

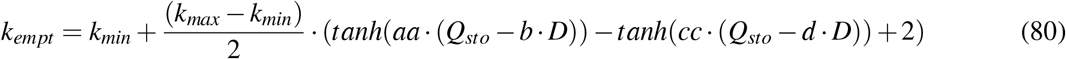

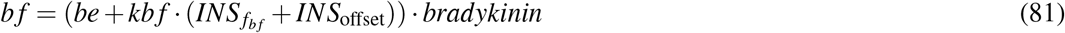

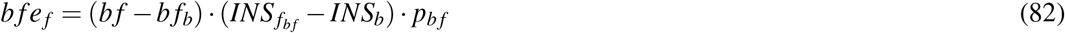

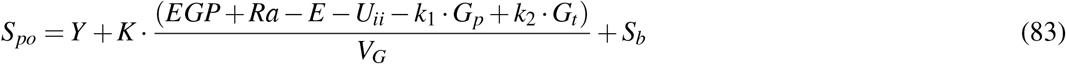

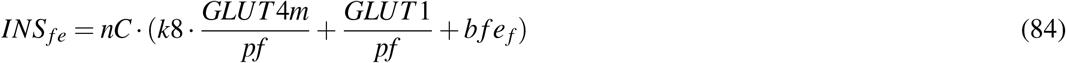

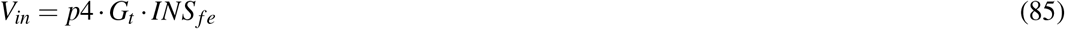

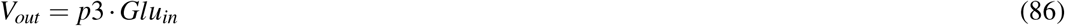

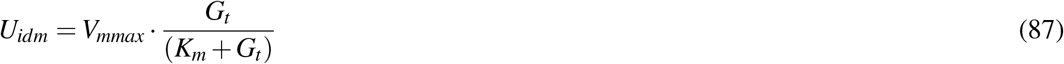

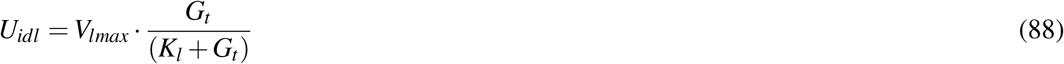

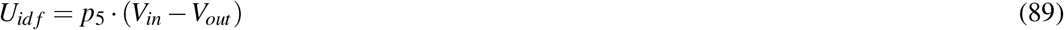

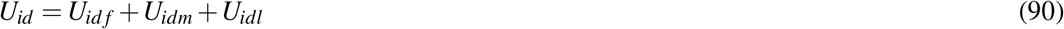

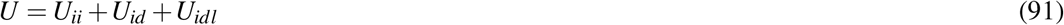

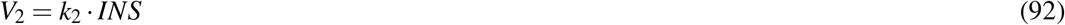

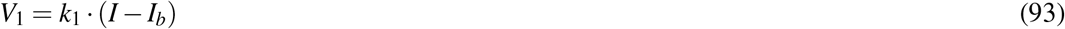

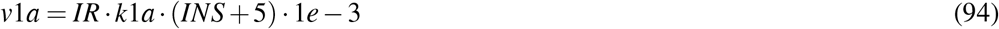

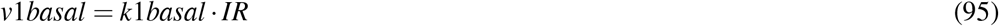

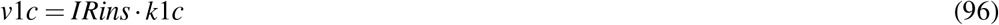

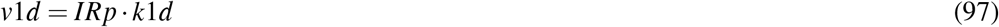

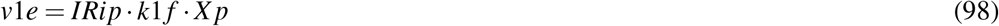

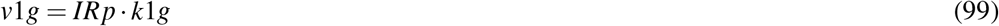

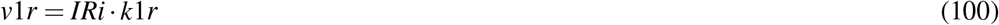

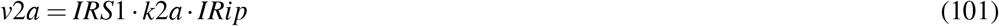

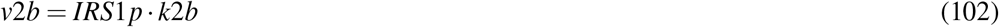

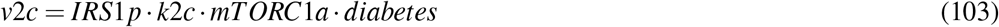

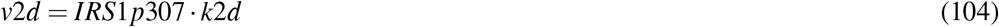

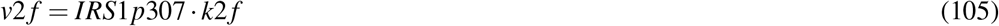

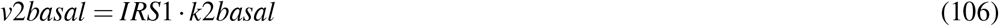

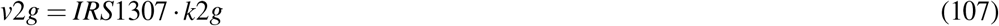

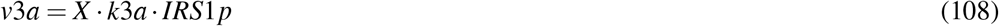

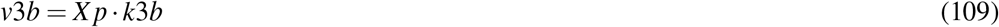

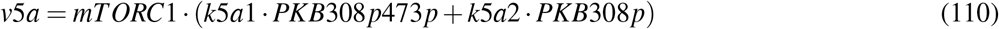

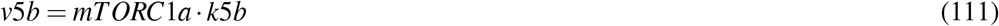

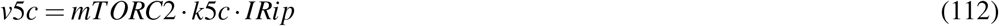

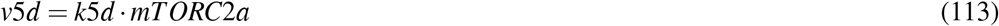

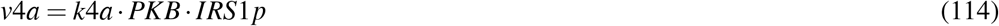

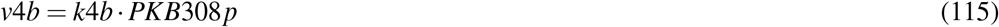

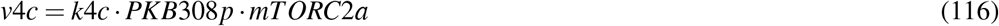

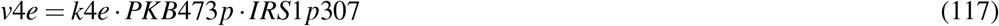

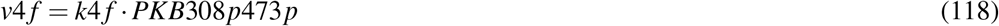

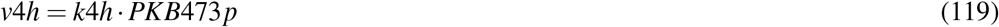

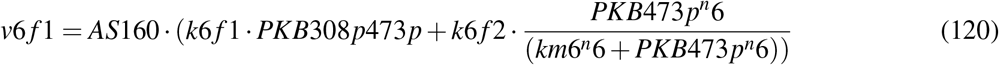

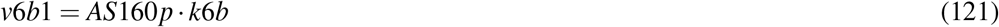

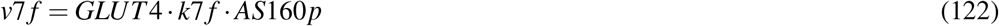

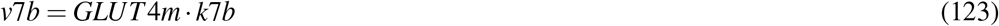

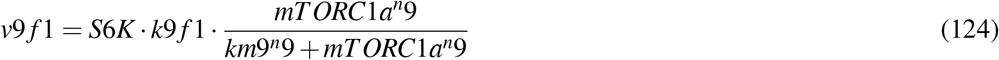

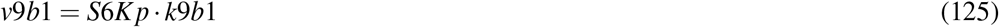

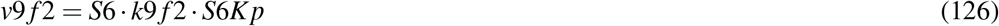

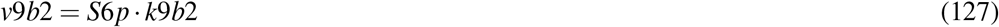

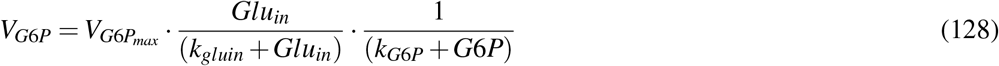

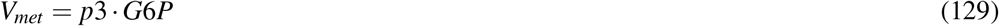

---

**Figure S1:**
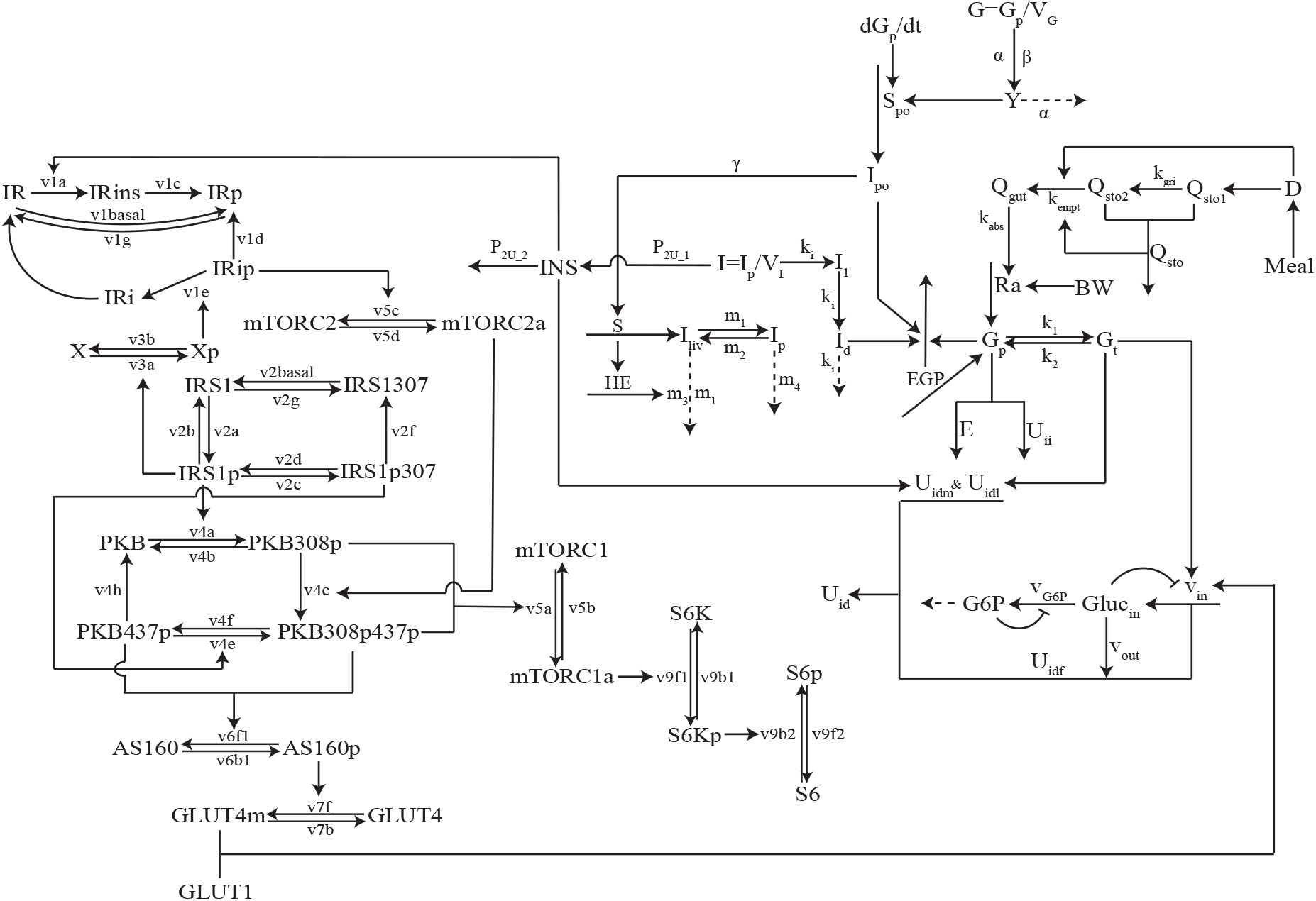
Interaction graph for model M2b.

**Figure S2:**
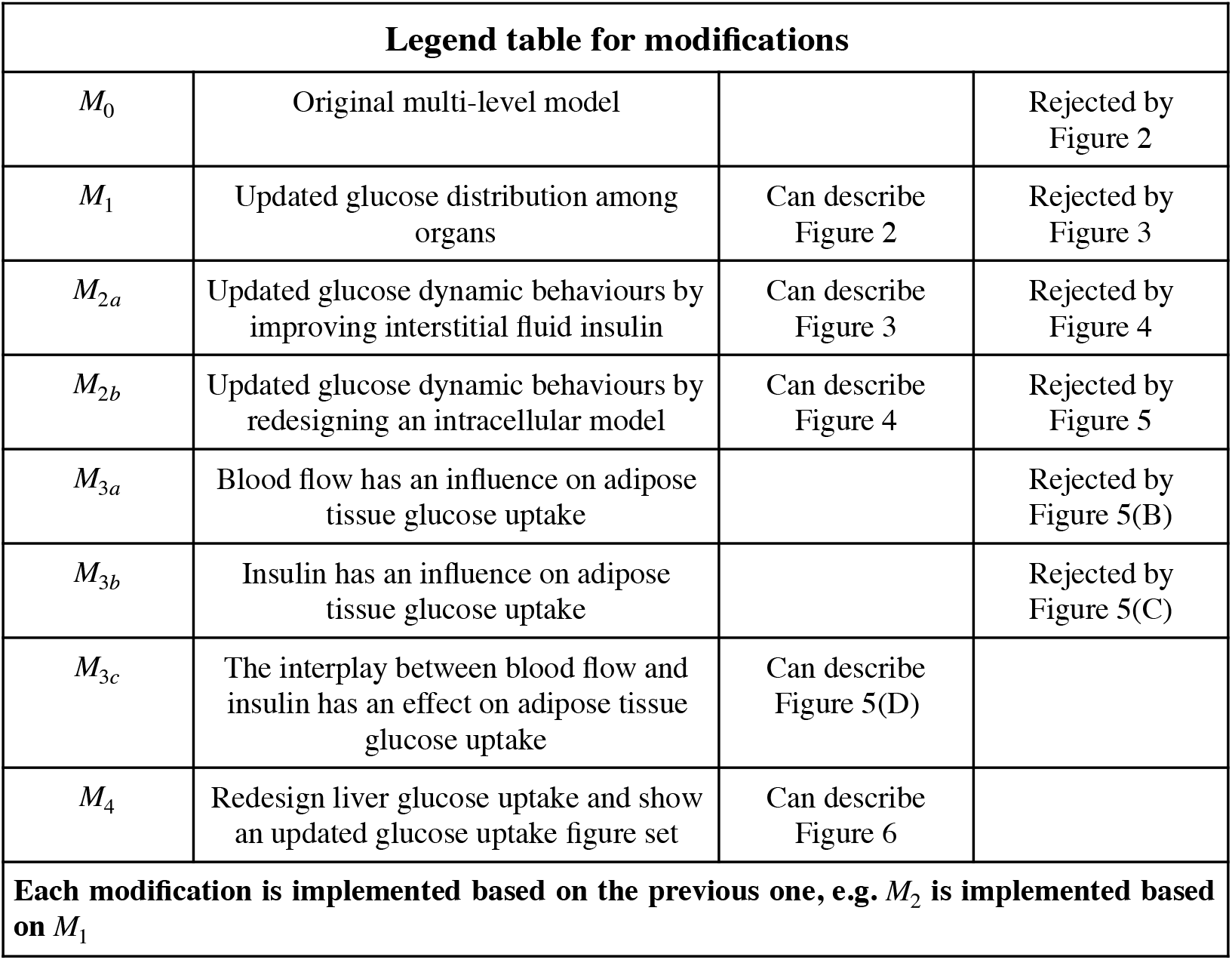
All evaluated models.

